# Pervasive but lagged responses in the composition of small mammal communities to a century of climate change

**DOI:** 10.1101/2024.04.24.590797

**Authors:** Ethan Abercrombie, Jonathan A. Myers, Adam B. Smith

## Abstract

Climate change has widespread effects on the distribution, abundance, and behavior of species around the world, leading to the reshuffling of ecological communities. However, it remains unclear whether individual species’ range shifts scale up to result in communities whose rate of change lag, lead, or track the rate of climate change. We capitalized on a century-old data set originally collected by Joseph Grinnell and his students, plus modern resurveys, to measure long-term compositional responses of small mammal communities to climate change in historical and modern eras across three regions in the Sierra Nevada of California (Lassen, Yosemite, and Sequoia and Kings Canyon National Parks). Across this period, mean annual temperature in each region increased and mean annual precipitation decreased. We tested whether small mammal communities have shifted their composition in favor of species more adapted to hot and dry conditions, processes known as thermophilization and mesophilization respectively. We found positive thermophilization rates (communities composed of more warm-adapted species) in all three regions and negative mesophilization rates (communities composed of dry-adapted species) in one of the three regions. Importantly, thermophilization and mesophilization rates lagged behind corresponding rates of climate change on average by 0.90-1.2 °C and 26.5-111.0 mm, but the magnitude of lags was unrelated to region, amount of climate change, or their interaction. Previous work demonstrated high intraspecific variability in range shifts across the three regions in our study. Our results suggest that the net effects of climate change will be directional at the scale of the ecological community, despite variability in individual species responses to environmental change and the varied mechanisms that govern them. Communities, like many individual species, may already be out of equilibrium with ambient climate.

## Introduction

Climate change has widespread effects on the distribution (Chen et al. 2011), abundance (Bowler et al. 2017), and behavior (Loe et al. 2016) of species around the world, leading to the reshuffling of ecological communities (Williams and Jackson 2007). The effects of climate change are particularly pronounced in montane ecosystems where compressed environmental gradients lead to rapid turnover in biodiversity across elevation (Sanders and Rahbek 2012, Graham et al. 2014). One prominent hypothesis to explain biodiversity responses to climate change is that species will track their preferred climate conditions by migrating to new areas with suitable climates (Colwell et al. 2008, Pecl et al. 2017). This climate-tracking hypothesis makes two predictions: 1) individual species exhibit range shifts concomitant with their climate preferences, and 2) ecological communities exhibit directional changes in composition, with increases in the relative abundances of species best-adapted to novel climate conditions at a site. In montane ecosystems, it is generally thought that species will move upslope to track preferred temperature regimes (Thuiller 2007, Feeley et al. 2012, Lenoir and Svenning 2015). However, there is mixed support for upslope climate-tracking in montane ecosystems. While many species have moved to higher elevation in recent decades (Chen et al. 2011, Mamantov et al. 2021), a considerable number have not shifted their elevational distributions or have migrated downslope (Rubenstein et al. 2023). Given the variable responses of individual species to climate change, it remains unclear whether individual species’ range shifts scale up to result in directional changes in the composition of entire communities.

Several processes can lead to directional or non-directional responses of montane communities to climate change. First, communities can display directional change despite idiosyncratic range shifts if overall effects of climate on the local community are strong (Devictor et al. 2012, Fadrique et al. 2018). For example, many communities are shifting in composition towards more warm-adapted and dry-adapted species, a process known as thermophilization and mesophilization respectively (Feeley et al. 2012, Duque et al. 2015, Ramón-Martínez and Seoane 2024). These directional changes can result from upslope migration (Morueta-Holme et al. 2015), and selective extirpation of less-adapted species (Freeman et al. 2018). Second, communities can display non-directional change due to variable effects of dispersal limitation or biotic interactions on species range shifts. Dispersal limitation may limit the migration of species, leading to elevational distributions that are decoupled from preferred climate (Wen et al. 2022). Likewise, elevational changes in the strength of local biotic interactions such as competition can result in downslope, upslope, or no change in species ranges (HilleRisLambers et al. 2013, Alexander et al. 2015). If the majority of species respond to different environmental and biotic factors, the signature of climate change at the community-level may be obscured.

Three key factors have generally limited our understanding of how species’ range shifts scale up to community-level outcomes. First, there is overemphasis on the role of temperature in driving species distributions, when other climatic factors such as precipitation may be important (Tingley et al. 2009, Ackerly et al. 2010). For example, while climate warming is expected to increase upslope migration of species, precipitation and water balance can facilitate downward migration (Lenoir et al. 2010). Second, long-term community assessments are rare. Despite the importance of long-term studies in furthering theory and understanding biodiversity responses to global change (Franklin et al. 1990, Kuebbing et al. 2018), the duration of most ecological research is around five years (Estes et al. 2018). Century-long studies are exceedingly uncommon but offer some of the best evidence for climate change’s impact on biodiversity (Tingley et al. 2009, Morueta-Holme et al. 2015). Lastly, most research has focused on single elevational transects, although species have been shown to respond differently throughout their range (Tingley et al. 2012, Rapacciuolo et al. 2014; Rowe et al. 2015).

In this study, we tested whether the composition of small-mammal communities has changed directionally over a century of climate change in three montane regions, and whether their rate of change tracks the rate of climate change. In a previous study, Rowe et al. (2015; also cf. Moritz et al. 2008) estimated the historical and contemporary elevational ranges for small mammals across the Sierra Nevada of California, USA, using species survey data from sites originally sampled by Joseph Grinnell and his students in the early 1900s (Grinnell and Storer 1924, Grinnell et al. 1930, Sumner and Dixon 1953), and resurveys of the same sites from 2003 to 2011. Notably, the detail with which Grinnell recorded observations of animals and the modern study design allowed the application of occupancy-detection modeling (MacKenzie et al. 2018) of data from both periods to remove bias from estimates of elevational ranges. Rowe et al. (2015) found no single species shifted their range in the same way across three different regions (Lassen Volcanic National Park and National Forest, Yosemite National Park, and Sequoia and Kings Canyon National Parks). We capitalize on this nearly century-old data set, to explicitly test whether the net effects of climate change on small mammal communities are directional in the face of high variability in individual species’ range shifts. We incorporate two aspects of climate, temperature and precipitation, to understand how ecological communities have changed over the past one hundred years. Mean annual temperature has increased, and mean annual precipitation has decreased, in all three regions between historical and contemporary surveys (Fig. 1). Specifically, we test the prediction that small mammal communities are comprised of more warm-adapted and dry-adapted species after a century of climate warming and drying, and that the rate of community-level change in thermophilization and mesophilization rates has tracked the rate of climate change in temperature and precipitation.

**Figure 1:**
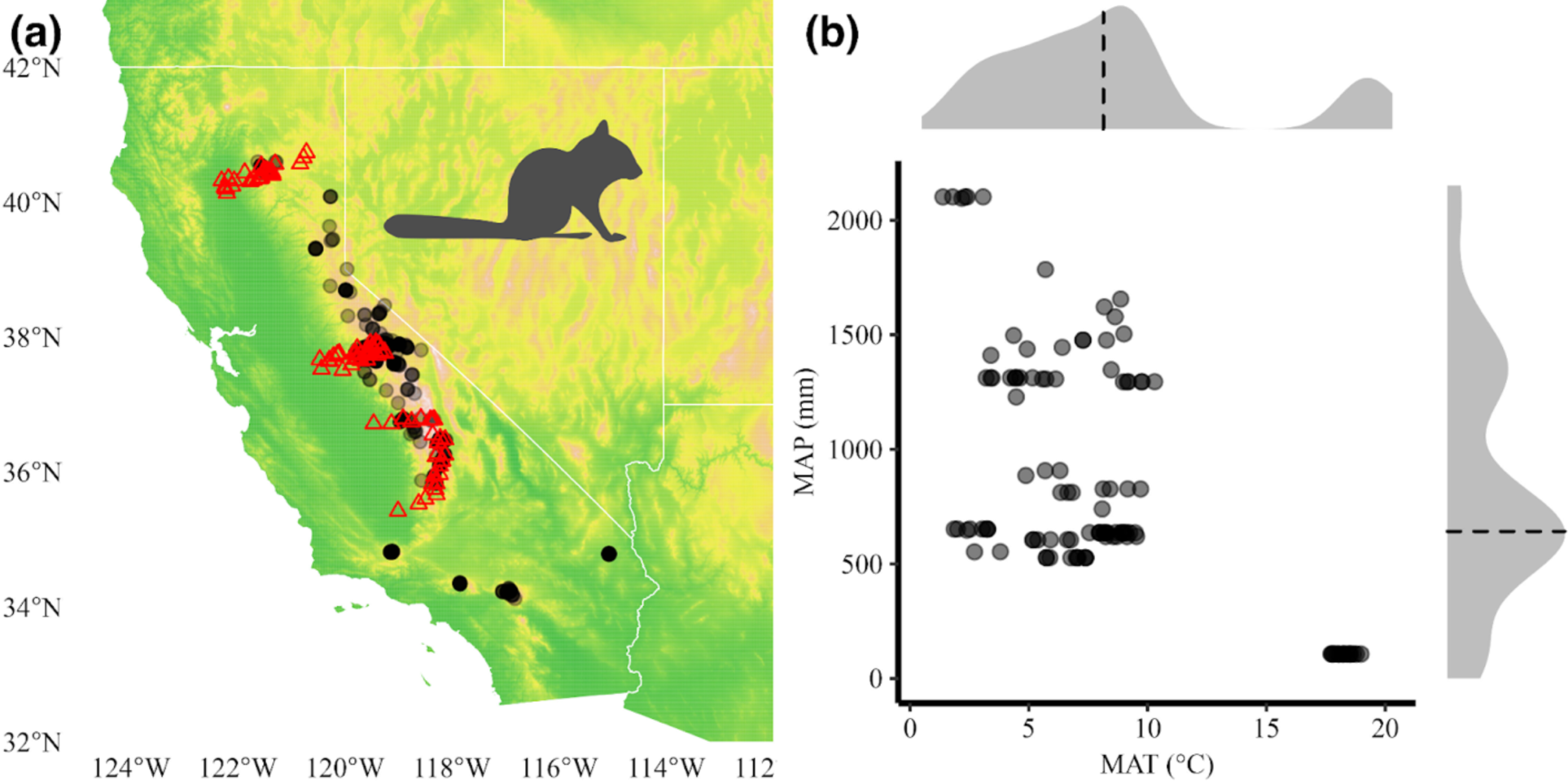
Methodology for calculating the community temperature index (CTI) and community precipitation index (CPI). a) For each species (data for *Tamius speciosus* depicted here) we collected occurrence records from across the entire species range. Black circles represent occurrence records. Red triangles represent field sites. b) We measured each species’ preferred temperature and precipitation by taking the median mean annual temperature (MAT; dashed vertical line) and mean annual precipitation (MAP; dashed horizontal line), respectively, from the location of each occurrence (31-year averages, ± 15 years from the year of collection). The CTI and CPI of each site, in each era, were measured as the median of all preferred temperatures and precipitation regimes for each species present. Color shading in panel a) shows elevation (green = lower elevations; yellow / orange = higher elevations).

## Methods

We evaluated compositional responses of small-mammal communities using community-wide climate indices commonly used to study how ecological communities respond to climate change (Devictor et al. 2012, Feeley et al. 2013). To generate these indices, we first used historical and modern species’ elevational range data from three elevational transects to determine species composition at a site. We then averaged the preferred temperature and precipitation values of each species present at a site, estimated using the location of range-wide occurrence records for each species, to calculate a community temperature index (CTI) and community precipitation index (CPI) for historical and modern eras. We detail each of the steps below (Fig. 1).

### Small mammal range data

Species’ elevational range data for the 34 nonvolant small-mammal species in this study (Appendix S1: Table S1), namely rodents and lagomorphs, come from Moritz et al. (2008) and Rowe et al. (2015), and were generated as part of the Grinnell Resurvey Project (https://mvz.berkeley.edu/Grinnell/index.html) at the University of California’s Museum of Vertebrate Zoology (https://mvz.berkeley.edu). Joseph Grinnell and his students surveyed California’s small mammal and bird diversity, including sites across three transects spanning the elevational gradient in the Sierra Nevada (Grinnell and Storer 1924, Grinnell et al. 1930, Sumner and Dixon 1953). Oak woodland and chaparral dominate at low elevations, giving way to mixed coniferous forest at intermediate elevations, and alpine conditions at high elevations. Small-mammal composition was recorded from 1911 to 1934 and include 34 sites within and around Lassen National Park and National Forest in the southern Cascade range (northern region, 91-2487 m), 47 sites from Yosemite National Park (central region, 118-3419 m), and 53 sites from Sequoia and Kings Canyon National Parks (southern region, 63-3292 m). Moritz et al. (2008) and Rowe et al. (2015) resurveyed these transects between 2003 and 2010, including 38 sites in Lassen, 81 sites in Yosemite, and 47 sites in Sequoia and Kings Canyon. Using these data, Moritz et al. (2008) and Rowe et al. (2015) estimated the elevational range shifts (hereafter range shifts) of nonvolant, non-game small mammals across these three regions using occupancy-detection modeling to correct for the problem of non-detection (Moritz et al. 2008, Rowe et al. 2015). We used the modeled historical and modern elevational ranges of nonvolant mammals in northern, central, and southern regions.

### Species climate preferences

We estimated the thermal and precipitation preferences for the 34 small-mammal species using occurrence data downloaded from the Global Biodiversity Information Facility (data DOI: https://doi.org/10.15468/dl.7x7sex). Thermal and precipitation preferences were estimated across the entire range of each species using climate data at the location of each occurrence record. Previous research using these methods typically use 30-year climate normals when assigning climate values to each record (e.g., Fadrique et al. 2018). We improve upon these methods by considering the exact year each record was collected, plus preceding years, which allows us to account for changes in climate at any given location. We first filtered all occurrences to those recorded as “occurrence”, or those representing collected specimens, and collected between 1901 and 2021. All records without latitude and longitude were removed. To remove spurious records, we used species’ range maps from the IUCN (International Union for Conservation of Nature 2018). For a given species, specimens located outside a buffer of 240 km around its range map were excluded. We chose a 240 km buffer because visual inspection indicated it gave a balance of excluding outliers and including parts of distributions not reflected in range polygons. If coordinate uncertainty was not reported, occurrences were discarded unless the occurrence was found within the IUCN range map. All occurrences with coordinate uncertainty greater than 1000 m were excluded. Lastly, we excluded any occurrences with any remaining geospatial issues. Mean annual temperature (MAT) and mean annual precipitation (MAP) from 1901 to 2021 were extracted for each occurrence using ClimateNA software version 7.30 (Wang et al. 2016). The recorded temperature and precipitation for each occurrence was measured as the 31-year average (±15 years) around the year of collection. We included additional years before and after the date of collection to ensure a span of 31 years for records without 15 years of available climate data before and after the date of collection (e.g., for records collected <15 years after 1901 and <15 years before 2021, the first and last years in which weather data was available). Species’ thermal and precipitation preferences were measured as the median temperature and precipitation of the 31-year averages of each record (Fig. 1). The final data set used to measure species’ thermal and precipitation preferences included 56,041 occurrences across all species. Preferences varied considerably across species (Appendix S1: Table S2).

### Community climate indices

For each historical site originally surveyed by Grinnell, we created a species-by-site matrix using presence/absence data for historical and modern eras. A species was deemed present if the site’s elevation overlapped with the species’ elevational range for each respective era. We measured the historical and contemporary community temperature index (CTI) and community precipitation index (CPI) at each of these sites as the median of species’ thermal and precipitation preferences, respectively, for all species present. Because our analysis utilized species presence/absence data (Rowe et al. 2015), changes in the community climate indices at each site are the result of changes in species composition. Thermophilization and mesophilization rates were calculated for each region separately by dividing the net change in the CTI and CPI by the average number of years spanning historical and modern surveys (northern = 81 years, central = 84 years, southern = 96 years). We used ClimateNA (Wang et al. 2016) and the same time spans to calculate changes in mean annual temperature (MAT) and mean annual precipitation (MAP) at each site (± 15-years from historical and modern censuses, northern = 1926-2007, central = 1916-2010, southern = 1913-2009). We included additional years above or below historical and modern censuses to ensure a span of 31 years for censuses without 15 years of available climate data before and after its date of occurrence.

## Statistical analyses

All analyses were performed in R version 4.3.1 (R Core Team 2023). Data tidying and formatting were performed using tidyverse packages (Wickham et al. 2019). All spatial procedures used with occurrence data utilized functions from the sf package (Pebesma 2023). We used a one-tailed t-test to test whether communities in each region have positive thermophilization and negative mesophilization rates (we used a one-tailed test specifically to evaluate whether communities have tracked the directionality of climate change toward warmer and drier conditions). To test for differences in thermophilization and mesophilization rates between regions, we used a one-way ANOVA. We evaluated whether small mammal communities were tracking climate by comparing changes in site MAT and MAP with changes in the CTI and CPI, respectively, using a Welch’s unequal variance t-test for each region. To evaluate whether the magnitude of site warming had an effect on the magnitude of change in the CTI, we used an ANOVA with CTI change as the response variable, and MAT change, region, and the interaction between MAT change and region as predictor variables. To evaluate whether the magnitude of site drying had an effect on the magnitude of change in the CPI, we used an ANOVA with CPI change as the response variable, and MAP change, region, and the interaction between MAP change and region as response variables.

## Results

Over the past ∼100 years mean annual temperatures have risen in all three regions (Fig. 2). On average, northern sites have risen 1.1°C, central sites 1.4°C, and southern sites 1.5°C. Temperatures rose more at higher elevations than lower elevations in central and southern sites. Mean annual precipitation has decreased 79 mm on average in northern sites, 87 mm in central sites, and 120 mm in southern sites (Fig. 2). An important exception to these patterns is low-elevation northern sites that have higher precipitation in the modern era. All sites showed elevation-dependent drying, with higher elevations experiencing the largest decreases in precipitation.

**Figure 2:**
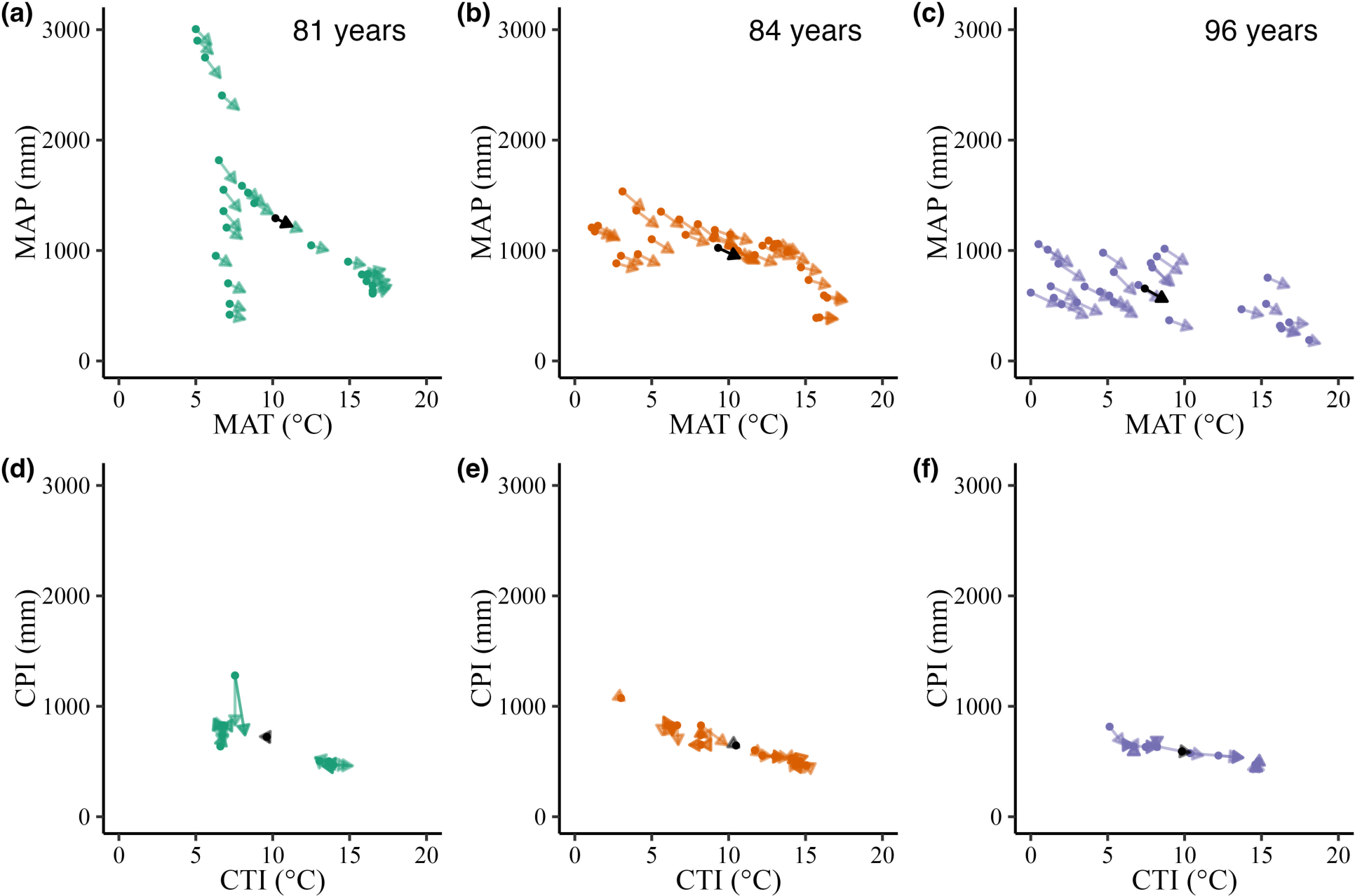
Climate vectors for site mean annual temperature (MAT) and mean annual precipitation (MAP) (a,b,c), and the community temperature index (CTI) and community precipitation index (d,e,f). a,d) Northern sites (Lassen National Park and National Forest; 81 years: 1926-2007), b,e) central sites (Yosemite National Park; 84 years: 1916-2010), c,f) southern sites (Sequoia and Kings Canyon National Parks; 96 years: 1913-2009). Points represent historical values for each site while arrow tips represent contemporary climate. Black points represent the mean historical value while black arrow tips represent mean contemporary values. Nearly all climate vectors indicate sites have become hotter and drier.

### Community responses to climate warming and drying

We observed positive and significant thermophilization rates (change in community composition towards species preferring warmer conditions) across all three regions (one-sided t-test, Fig. 3). In the northern region, the mean thermophilization rate across sites was significantly greater than zero (2.42 x 10^-3^ °C yr^−1^, t = 2.41, P = 0.01) and positive in 80% of sites (Fig. 3, Appendix S1: Figure S1). In the central region, the mean thermophilization rate was significantly greater than zero (2.34 x 10^-3^ °C yr^-1^, t = 2.04, P = 0.02) and positive in 51% of sites. In the southern region, the mean thermophilization rate was significantly greater than zero (5.66 x 10^-3^ °C yr^-1^, t = 3.17, P < 0.01) and positive in 63% of sites. We observed the greatest thermophilization rates at mid-elevations in southern sites (Fig. 3). Mean thermophilization rates did not differ significantly between sites (ANOVA, F2,84 = 1.92, P = 0.15).

**Figure 3:**
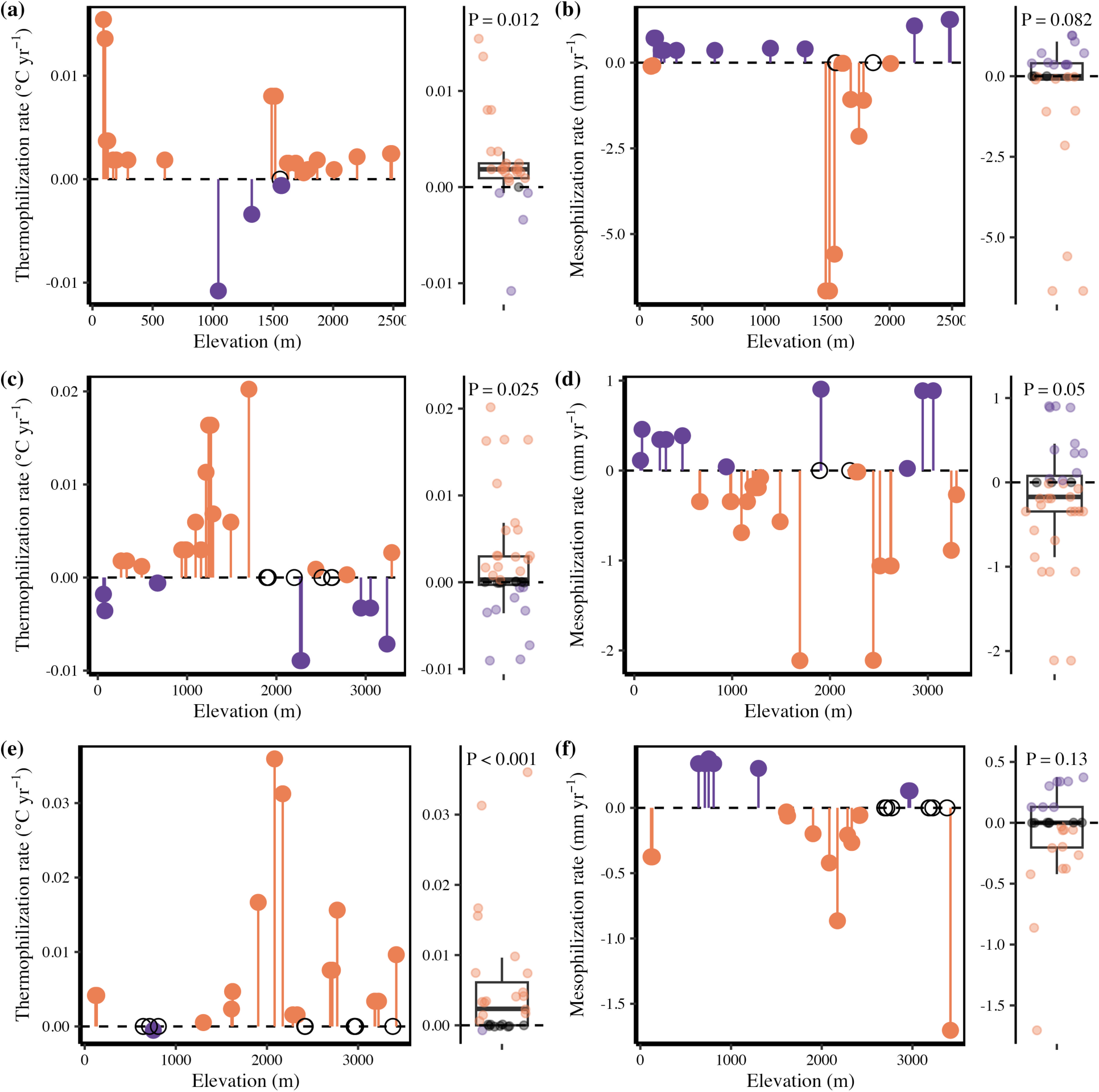
Thermophilization rates (°C yr^>−1^) and mesophilization rates (mm yr^−1^) of small mammal communities at sampled sites in the three study regions across elevation. a, b) Northern region (Lassen National Park); e, f) Central region (Yosemite National Park); C, D) Southern region (Sequoia and Kings Canyon National Parks). Box plots show region-wide trends in thermophilization and mesophilization rates and P-values from one-tail t-tests. Each point in the boxplot corresponds to a point in the graph to the left of it. Colors represent communities displaying warming and/or drying (orange), cooling and/or wetting (purple), or no change in community climate indices (black open circle). Communities with thermophilization and mesophilization rates of 0 represent communities whose composition has not changed since original surveys.

We observed negative mesophilization rates (change in community composition towards species preferring drier conditions) across all three regions, but only the central region was significant (one-sided t-test, Fig. 3). In the northern region, the mean mesophilization rate did not differ significantly from zero (-0.65 mm yr^-1^, t = -1.43, P = 0.08) and negative in 50% of sites (Fig. 3, Appendix S1: Figure S1a). In the central region, the mean mesophilization rate was significantly less than zero (-0.20 mm yr^-1^, t = -1.69, P < 0.05) and negative in 60% of sites. In the southern region, the mean mesophilization rate did not differ significantly from zero (-0.09 mm yr^-1^, t = -1.15, P = 0.13) and was negative in 44% of sites. Mean mesophilization rates did not differ significantly between sites (ANOVA, F2,84 = 1.31, P = 0.27).

### Lagged responses of community composition to climate change

We observed lags in the CTI compared with changes in site mean annual temperature (MAT) in all three regions (Fig. 4; Appendix S1 Figure S2). Among northern sites, the mean increase in site MAT was 1.1°C while the mean increase in the CTI was 0.20°C (t = 10.76, P < 0.001). Among central sites, the mean increase in site MAT was 1.4°C while the mean increase in the CTI was 0.20°C (t = 12.39, P < 0.001). Among southern sites, the mean increase in site MAT was 1.5°C while the mean increase in the CTI was 0.54°C (t = 5.19, P < 0.001). We observed no significant effect of region (ANOVA, F2,81 = 1.74, P = 0.18), MAT change (F1,81 = 0.89, P = 0.35), or the interaction between MAT change and region on the change in the CTI (F2,81 = 0.42, P = 0.66).

**Figure 4:**
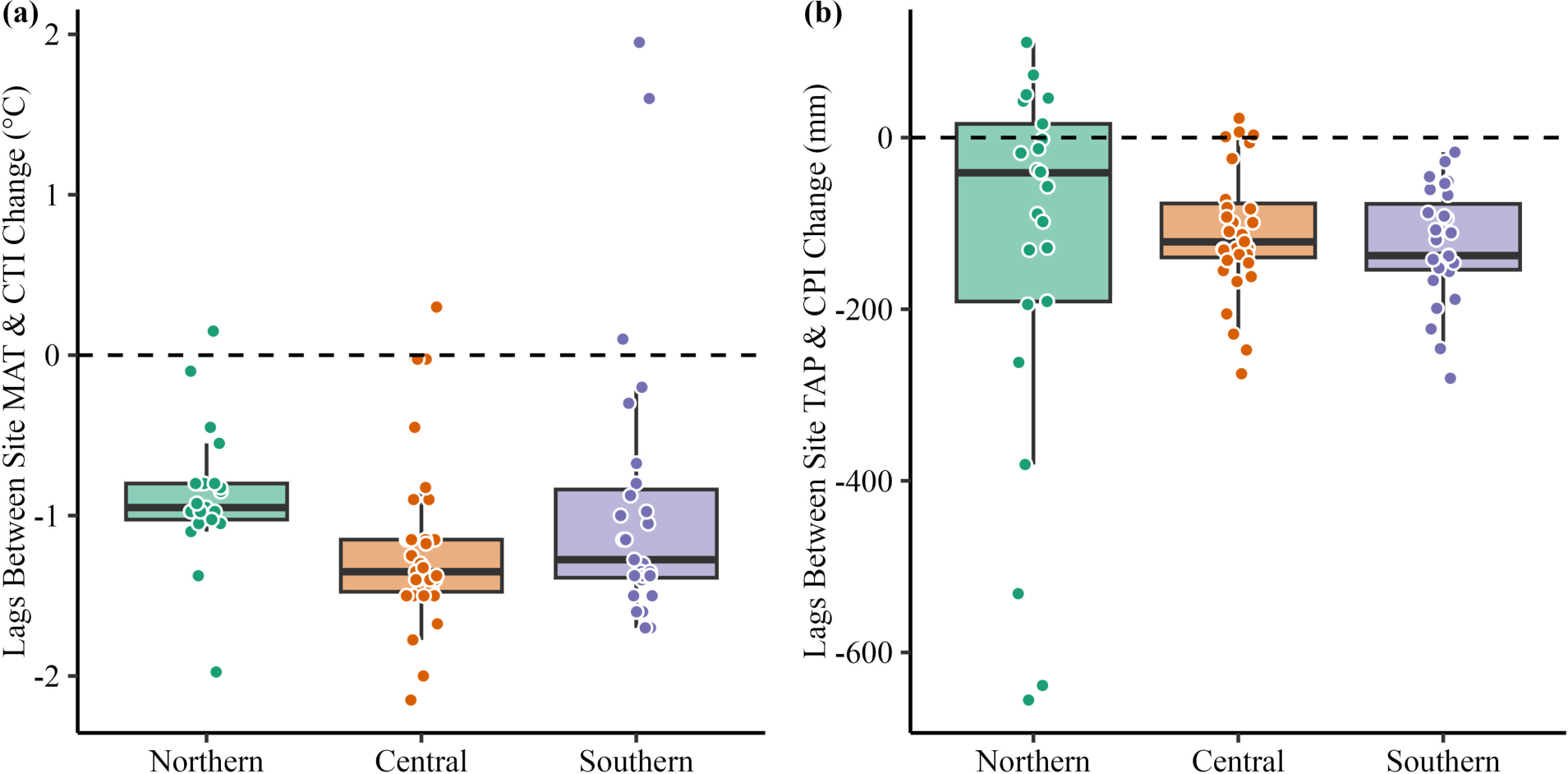
Disequilibrium in the magnitude of change between site climate and compositional change. a) The difference between site temperature change (°C) and change in the CTI (°C). b) The difference between site precipitation change (mm) and change in the CPI (mm).

We observed lags in the CPI compared with changes in site mean annual precipitation (MAP) in central and southern sites, but not northern sites (Fig. 4; Appendix S1 Figure S2). Among northern sites, the mean decrease in MAP was 79 mm while the mean decrease in the CPI was 52.5 mm (t = -0.50, P = 0.62). Among central sites, the mean decrease in MAP was 87 mm while the mean decrease in the CPI was 17.0 mm (t = -6.14, P < 0.001). Among southern sites, the mean decrease in MAP was 120 mm while the mean decrease in the CPI was 9.0 mm (t = -9.16, P < 0.001). We observed no significant effect of region (ANOVA, F2,81 = 1.90, P = 0.16), MAP change (F1,81 = 2.65, P = 0.11), or the interaction between MAP change and region on the change in the CPI (F2,81 = 0.31, P = 0.74).

## Discussion

The composition of small-mammal communities across montane California are shifting in favor of more warm- and dry-adapted species but not at a pace required to track climate change. This consistent directional change in composition occurred despite high levels of intraspecific heterogeneity in responses to climate change across the three regions (Rowe et al. 2015). MAT increased at all sites in our study (Fig. 2). Except for the warmest low-elevation northern sites, MAP decreased (Fig. 2). In line with a recent analysis showing elevational-dependent warming is a common phenomenon worldwide (Pepin et al. 2015), the magnitude of the increase in MAT in central and southern sites increased with elevation.

Species range shifts in response to recent climate change has led to positive thermophilization rates across all three regions, and negative mesophilization rates in two of the three regions. Importantly, changes in the composition of small mammal communities are lagging behind changes in climate (Fig. 4). While we acknowledge the difficulty for any one theory to predict the direction and magnitude of species’ responses to climate change, our findings provide evidence for directional responses of communities to climate change concomitant with ongoing changes in temperature and precipitation. Our study also demonstrates that communities, like many individual species, may already be out of equilibrium with ambient climate.

Compared to the past, small mammal communities across the Sierra Nevada are comprised of more warm-adapted and dry-adapted species. Thermophilization rates were positive across all three regions, and effect sizes were comparable to previous research in other ecosystems on birds (Devictor et al. 2012), butterflies (Devictor et al. 2012), and plants (De Frenne et al. 2013, Fadrique et al. 2018). Furthermore, mesophilization rates were negative among central sites. These results suggest decreases in precipitation across the Sierra Nevada have also contributed to directional shifts in community composition. Two mechanisms can contribute to positive thermophilization and negative mesophilization rates among our sites: range expansions and range contractions. In recent years, high-elevation (cool/wet-adapted) species including the alpine chipmunk (*Tamius alpinus*) and the Belding’s ground squirrel (*Urocitellus beldingi*) have shown range contractions at the lower edge of their ranges (Rubidge et al. 2011, Morelli et al. 2012), while low-elevation (warm/dry-adapted) species like the pinyon mouse (*Peromyscus truei*) and the Californian pocket mouse (*Chaetodipus californicus*) show range expansions at their upper edges (Fig. 1, Rowe et al. 2015). Both mechanisms, the expansion of low-elevation species and the contraction of high-elevation species, contribute simultaneously to the high thermophilization rates and negative mesophilization rates at mid-elevations (Fig. 3).

Although we observed region-wide trends of thermophilization and mesophilization in our study, both of these rates were highly variable across sites (Fig. 3). Biotic interactions, vegetation dynamics, and behavioral buffering are three mechanisms that can increase variation in thermophilization and mesophilization rates. For example, low- and mid-elevation species in Yosemite expanded their ranges by tracking suitable vegetation (Santos et al. 2015). Slow rates of thermophilization and mesophilization might be due to the inability for mammals to shift their ranges because of a lack of suitable vegetation. Furthermore, low rates of thermophilization and mesophilization might be due to the behavioral buffering capacity of mammal species at a site, delaying extirpation. For instance, small-mammal communities in California’s Mojave Desert were found to be largely stable over the past century (with only moderate turnover, ∼2 species), despite increases in temperature, because of low physiological exposure to climate change through burrowing (Riddell et al. 2021).

If mammals across elevation can behaviorally thermoregulate through microhabitat selection, species may not shift their ranges and thermophilization and mesophilization rates will be low. Considering how species range shifts are not the same across the three regions in our study (Rowe et al. 2015), it is surprising that thermophilization rates and mesophilization rates did not differ between regions. Instead, we find the community climate indices have changed similarly despite differences in the rate of climate change.

Importantly, thermophilization rates lag behind temperature changes in all regions, and negative mesophilization rates lag behind precipitation change in central and southern regions (Fig. 4). It is important to note that despite the nonsignificant difference between changes in site MAP and the CPI in the northern region, the majority of sites are still lagging behind climate change (Fig 4). While differences between thermophilization and mesophilization rates and compositional change can be explained through microhabitat selection, they can also occur from the inability for species to respond to climate change fast enough. If small mammal communities across the Sierra Nevada cannot keep pace with climate change, they are at risk of population collapse.

Our analyses highlight the importance of biological scale in studying the effects of climate change on biodiversity. Previous studies of species-level responses to climate change in the Sierra Nevada suggest no universal explanation for species’ range shifts (Rowe et al. 2015, Santos et al. 2015). We show that despite heterogeneous range shifts in small mammals across montane California, communities are displaying the signatures of climate warming and drying. Our findings demonstrate how the net effects of climate change can be directional at the scale of the ecological community, despite variability in individual species responses to environmental change and the varied mechanisms that govern them.

The lags between community-wide responses and climate change indicate that these communities are already out of equilibrium with ambient climate change. Compared to future forecasts (Cayan et al. 2008) only moderate warming and drying has occurred in the Sierra Nevada. Hence, we should expect that the degree of disequilibrium between these communities and their environments will only increase into the future. In an influential study, Colwell et al. (2008) draw attention to “lowland biotic attrition” in the tropics, which they expect to occur because the warmest places in those regions will become intolerable warmer, while there are no nearby sources of species that can tolerate these extended conditions. Analogously, we suggest a similar dynamic, but not necessarily restricted to the lowlands of tropical ecosystems. Rather, our work demonstrates that lags between climate and communities can occur across elevations. Even if the species pool of a particular location contains species that could reside there, the accelerating pace of climate change may disallow them from shifting their ranges to match climate. As a result, entire elevational gradients may eventually suffer from a low-grade biotic attrition.

## Acknowledgments

We thank members of the Smith Lab, Myers Lab, and Tello Lab for helpful discussions and feedback on the manuscript. We thank Kevin Rowe for sharing the results of the occupancy-detection models. EA was supported by a National Science Foundation Graduate Research Fellowship (DGE-2139839) and the George Hayward Plant Biology Graduate Fellowship at Washington University in St. Louis. This work was partially supported by the Alan Graham Fund in Global Change to ABS.

