## Appendix S1 for "Pervasive but lagged responses in the composition of small mammal communities to a century of climate change"

Table S1: Historical and modern elevational ranges of the small mammals included in our analyses in Lassen, Yosemite, and Sequoia and Kings Canyon National Parks.

| **Era** | **Species** | **Lassen (Northern)** | **Yosemite (Central)** | **Sequoia & Kings Canyon (Southern)** |
|---|---|---|---|---|
| Historical | *Callospermophilus lateralis* | 1561-3124 | 1646-3200 | 2147-3474 |
| Modern | *Callospermophilus lateralis* | 1459-2491 | 1951-3255 | 2262-3454 |
| Historical | *Chaetodipus californicus* | X | 183-914 | 118-2147 |
| Modern | *Chaetodipus californicus* | X | 557-1701 | 231-2373 |
| Historical | *Clethrionomys californicus* | 1971-1971 | X | X |
| Modern | *Clethrionomys californicus* | 1459-2491 | X | X |
| Historical | *Dipodomys agilis* | X | X | 721-1860 |
| Modern | *Dipodomys agilis* | X | X | 810-2167 |
| Historical | *Dipodomys californicus* | 79-1051 | X | X |
| Modern | *Dipodomys californicus* | 207-1584 | X | X |
| Historical | *Dipodomys heermanni* | X | 52-975 | 118-636 |
| Modern | *Dipodomys heermanni* | X | 115-728 | X |
| Historical | *Marmota flaviventris* | 1561-1971 | 2469-3353 | 2268-3503 |
| Modern | *Marmota flaviventris* | 1561-2491 | 2469-3353 | 2268-3503 |
| Historical | *Microtus californicus* | 79-1335 | 52-1647 | 118-1261 |
| Modern | *Microtus californicus* | 80-1044 | 57-1646 | 583-999 |
| Historical | *Microtus longicaudus* | 1672-2462 | 583-3281 | 1529-3474 |
| Modern | *Microtus longicaudus* | 1468-2491 | 1227-3255 | 2167-3454 |
| Historical | *Microtus montanus* | 1335-1784 | 1211-3161 | 1984-3384 |
| Modern | *Microtus montanus* | 1468-1850 | 1205-3255 | 2167-3398 |
| Historical | *Neotoma cinerea* | 1478-2514 | 1803-3281 | 1529-3384 |
| Modern | *Neotoma cinerea* | 1680-1785 | 2383-2474 | 1507-3398 |
| Historical | *Neotoma fuscipes* | 79-1051 | X | X |
| Modern | *Neotoma fuscipes* | 111-1566 | X | X |
| sHistorical | *Neotoma macrotis* | X | 183-1647 | 118-2147 |
| Modern | *Neotoma macrotis* | X | 57-1701 | 231-2373 |
| Historical | *Ochotona princeps* | 1478-2514 | 2377-3871 | 2732-3384 |
| Modern | *Ochotona princeps* | 1478-2514 | 2377-3871 | 2732-3384 |
| Historical | *Otospermophilus beecheyi* | 79-1051 | 61-2632 | 118-2997 |
| Modern | *Otospermophilus beecheyi* | 111-1785 | 76-2784 | 138-2940 |
| Historical | *Peromyscus boylii* | 79-1051 | 183-2464 | 118-3147 |

| | | | | |
|---|---|---|---|---|
| Modern | *Peromyscus boylii* | 168-1044 | 57-2495 | 138-2282 |
| Historical | *Peromyscus californicus* | X | 251-1202 | 787-787 |
| Modern | *Peromyscus californicus* | X | 420-420 | 583-741 |
| Historical | *Peromyscus maniculatus* | 79-2514 | 52-3281 | 118-3384 |
| Modern | *Peromyscus maniculatus* | 92-2491 | 50-3255 | 135-3640 |
| Historical | *Peromyscus truei* | 79-1051 | 183-975 | 636-3147 |
| Modern | *Peromyscus truei* | 608-1459 | 557-1811 | 583-2940 |
| Historical | *Phenacomys intermedius* | X | 2386-3161 | 3460-3474 |
| Modern | *Phenacomys intermedius* | X | 2433-3255 | 3441-3441 |
| Historical | *Reithrodontomys megalotis* | 79-1478 | 52-1158 | 118-1860 |
| Modern | *Reithrodontomys megalotis* | 80-1044 | 57-1268 | 135-999 |
| Historical | *Sciurus griseus* | 103-1051 | 183-1951 | 787-2364 |
| Modern | *Sciurus griseus* | 103-1722 | 183-1689 | 1507-1614 |
| Historical | *Sorex monticolus* | X | 2176-3281 | 1529-3474 |
| Modern | *Sorex monticolus* | X | 1205-3255 | 1507-3441 |
| Historical | *Sorex ornatus* | 103-103 | 549-914 | 118-180 |
| Modern | *Sorex ornatus* | X | 57-569 | 231-1542 |
| Historical | *Sorex palustris* | 1583-2514 | 1647-3161 | 2314-3384 |
| Modern | *Sorex palustris* | 608-608 | 2153-3255 | 2990-3398 |
| Historical | *Sorex tenellus* | X | X | X |
| Modern | *Sorex tenellus* | X | 2936-3255 | 3240-3240 |
| Historical | *Sorex trowbridgii* | 1051-2061 | 1068-2286 | X |
| Modern | *Sorex trowbridgii* | 1459-2491 | 1129-2232 | 1507-2373 |
| Historical | *Sorex vagrans* | 1335-2514 | X | X |
| Modern | *Sorex vagrans* | 1044-2491 | X | X |
| Historical | *Tamias alpinus* | X | 2386-3353 | 2314-3503 |
| Modern | *Tamias alpinus* | X | 2883-3255 | 2785-3454 |
| Historical | *Tamias amoenus* | 1561-2514 | 2438-2865 | X |
| Modern | *Tamias amoenus* | 1559-2491 | 2474-2784 | X |
| Historical | *Tamias merriami* | X | 488-1524 | 636-2732 |
| Modern | *Tamias merriami* | X | 887-1268 | 999-2905 |
| Historical | *Tamias panamintinus* | X | X | 2732-2997 |
| Modern | *Tamias panamintinus* | X | X | 2772-3398 |
| Historical | *Tamias quadrimaculatus* | X | 1494-2210 | X |
| Modern | *Tamias quadrimaculatus* | X | 1646-2238 | X |
| Historical | *Tamias senex* | 1478-2462 | 1402-2743 | X |
| Modern | *Tamias senex* | 1459-2491 | 2383-2383 | X |
| Historical | *Tamias speciosus* | 1561-2514 | 1768-3281 | 1529-3384 |
| Modern | *Tamias speciosus* | 1783-2491 | 1881-3255 | 2167-3398 |
| Historical | *Tamiasciurus douglasii* | 886-2061 | 1229-3185 | 1592-3384 |
| Modern | *Tamiasciurus douglasii* | 886-2491 | 1229-3185 | 1592-3384 |

| | | | | |
|---|---|---|---|---|
| Historical | *Thomomys bottae* | 75-1335 | 57-1676 | 118-3384 |
| Modern | *Thomomys bottae* | 75-1335 | 57-1676 | 118-3384 |
| Historical | *Thomomys monticola* | 1561-2514 | 1905-3155 | X |
| Modern | *Thomomys monticola* | 1561-2514 | 1905-3155 | X |
| Historical | *Urocitellus beldingi* | 1485-1845 | 2286-3281 | 2761-3474 |
| Modern | *Urocitellus beldingi* | 1559-1628 | 2685-3255 | 3316-3454 |
| Historical | *Zapus princeps* | 1478-2462 | 1211-3281 | 1592-2657 |
| Modern | *Zapus princeps* | 1616-2276 | 1424-3255 | 2413-3240 |

*Notes:* X represents the absence of a species. All absent species were excluded from calculations of the community temperate index and community precipitation index.

Table S2. Species climate preferences as measured using occurrence records.

| Species | Preferred Precip (mm) | Min. Precip (mm) | Max. Precip (mm) | Preferred Temp ºC | Min. Temp ºC | Max. Temp ºC | # of clean records |
|---|---|---|---|---|---|---|---|
| *Callospermophilus lateralis* | 637.0 | 279 | 1467 | 6.60 | 2.6 | 9.4 | 975 |
| *Chaetodipus californicus* | 535.0 | 290 | 740 | 14.70 | 9.9 | 16.6 | 300 |
| *Dipodomys agilis* | 366.5 | 107 | 554 | 16.85 | 10.3 | 18.3 | 214 |
| *Dipodomys heermanni* | 295.0 | 207 | 453 | 16.30 | 12.6 | 17.6 | 261 |
| *Marmota flaviventris* | 626.0 | 395 | 1312 | 3.60 | 1.5 | 6.7 | 256 |
| *Microtus californicus* | 461.0 | 227 | 1217 | 14.90 | 8.6 | 17.5 | 787 |
| *Microtus longicaudus* | 652.0 | 519 | 1656 | 6.70 | 2.6 | 10.2 | 2425 |
| *Microtus montanus* | 649.5 | 463 | 1477 | 5.50 | 1.5 | 8.2 | 1590 |
| *Neotoma cinerea* | 597.0 | 161 | 1602 | 6.75 | 2.2 | 13.0 | 522 |
| *Neotoma fuscipes* | 929.0 | 357 | 1173 | 13.50 | 13.0 | 14.5 | 131 |
| *Neotoma macrotis* | 554.0 | 347 | 1217 | 14.60 | 8.5 | 17.3 | 249 |
| *Ochotona princeps* | 1479.0 | 375 | 1663 | 2.85 | 2.8 | 6.5 | 227 |
| *Otospermophilus beecheyi* | 531.0 | 261 | 1146 | 14.10 | 6.7 | 16.6 | 239 |
| *Peromyscus boylii* | 473.0 | 181 | 1350 | 13.80 | 9.0 | 18.3 | 3908 |
| *Peromyscus maniculatus* | 541.0 | 161 | 1466 | 8.50 | 3.0 | 17.5 | 22753 |
| *Peromyscus truei* | 454.0 | 107 | 1173 | 13.35 | 8.5 | 18.3 | 4325 |
| *Reithrodontomys megalotis* | 340.0 | 184 | 644 | 15.50 | 6.8 | 17.5 | 2409 |
| *Sciurus griseus* | 827.0 | 107 | 1543 | 12.20 | 8.5 | 18.3 | 76 |
| *Sorex monticolus* | 979.0 | 570 | 1602 | 6.05 | 2.3 | 8.2 | 3690 |
| *Sorex ornatus* | 1217.0 | 411 | 1217 | 14.90 | 14.9 | 16.8 | 105 |
| *Sorex palustris* | 830.0 | 541 | 1602 | 5.20 | 2.6 | 11.2 | 490 |
| *Sorex trowbridgii* | 1414.5 | 1332 | 1466 | 11.20 | 10.8 | 11.3 | 1004 |
| *Sorex vagrans* | 467.0 | 414 | 649 | 6.90 | 5.6 | 11.4 | 1625 |
| *Tamias alpinus* | 1075.0 | 647 | 2046 | 3.00 | 1.3 | 4.6 | 131 |
| *Tamias amoenus* | 826.0 | 405 | 1316 | 6.00 | 2.3 | 7.4 | 1011 |
| *Tamias merriami* | 359.0 | 107 | 929 | 16.10 | 9.1 | 18.3 | 121 |
| *Tamias quadrimaculatus* | 1512.0 | 1466 | 1622 | 9.90 | 8.8 | 11.2 | 36 |
| *Tamias senex* | 1613.0 | 1374 | 2102 | 8.20 | 2.3 | 10.2 | 125 |
| *Tamias speciosus* | 637.0 | 107 | 1785 | 8.20 | 2.3 | 18.3 | 462 |
| *Tamiasciurus douglasii* | 1231.0 | 527 | 1862 | 8.20 | 4.2 | 11.2 | 374 |

| | | | | | | | |
|---|---|---|---|---|---|---|---|
| *Thomomys bottae* | 466.0 | 107 | 1217 | 15.20 | 2.6 | 18.3 | 3714 |
| *Thomomys monticola* | 1477.0 | 1307 | 2096 | 5.00 | 2.2 | 8.2 | 192 |
| *Urocitellus beldingi* | 1327.0 | 1312 | 1742 | 4.20 | 3.0 | 6.7 | 138 |
| *Zapus princeps* | 1415.0 | 570 | 1670 | 8.20 | 3.6 | 11.8 | 1176 |

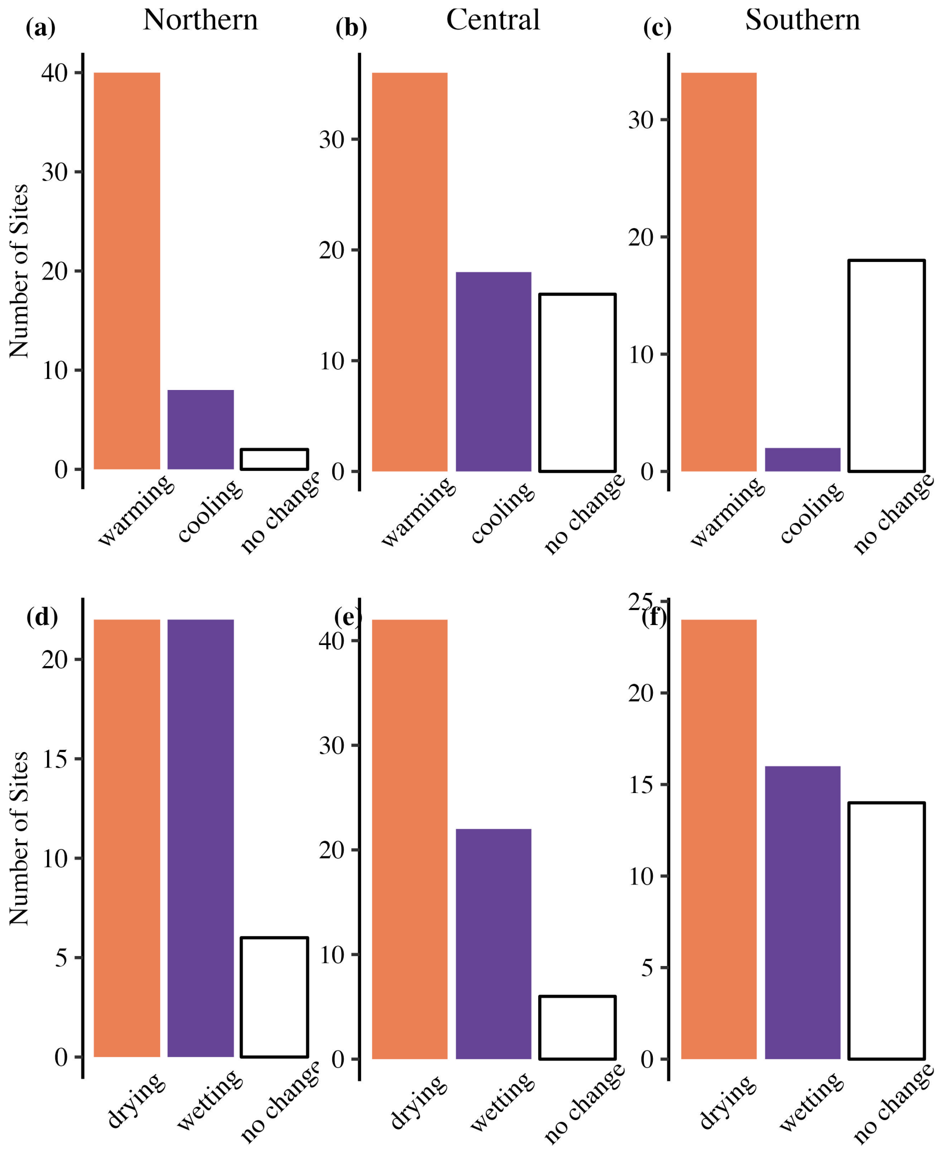


Figure S1: Frequency of sites with positive (warming), negative (cooling), and no change in thermophilization rates (a,b,c), and negative (drying), positive (wetting), and no change in mesophilization rates (d,e,f).

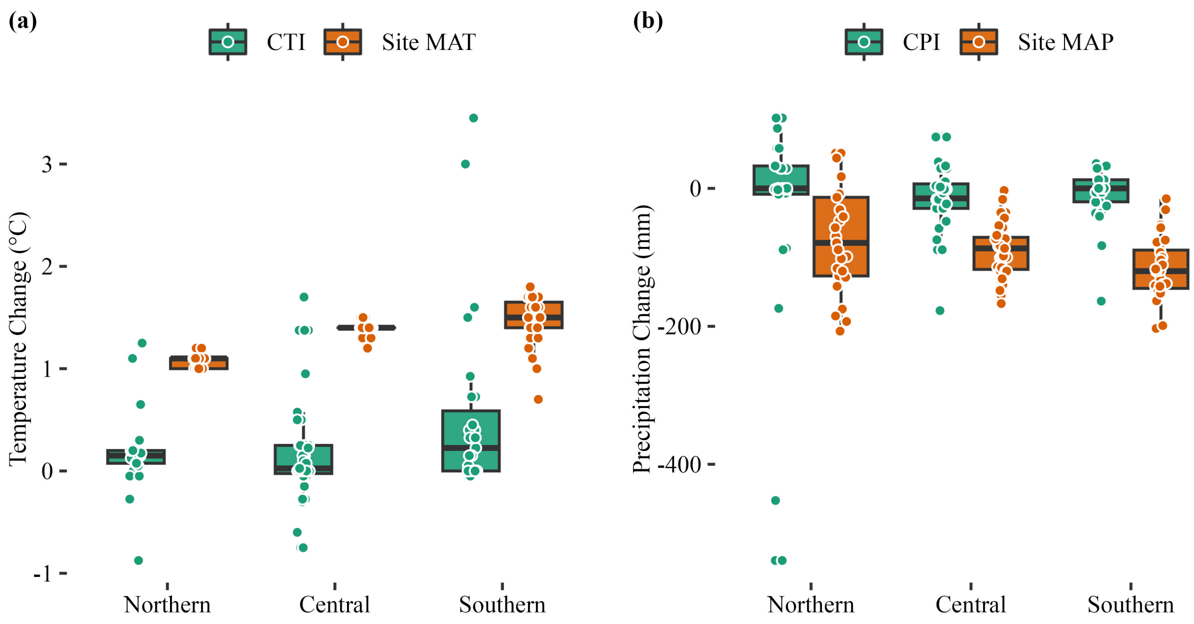


Figure S2: Changes in a) site temperature and CTI, and b) site precipitation and CPI. In most sites, changes in community composition lag behind changes in climate.
